# An alternative membrane topology permits lipid droplet localization of peroxisomal fatty acyl-CoA reductase 1

**DOI:** 10.1101/373530

**Authors:** Tarik Exner, Inés Romero-Brey, Eden Yifrach, Jhon Rivera-Monroy, Bianca Schrul, Christos C. Zouboulis, Wolfgang Stremmel, Masanori Honsho, Ralf Bartenschlager, Einat Zalckvar, Margarete Poppelreuther, Joachim Füllekrug

**Affiliations:** Molecular Cell Biology Laboratory Internal Medicine IV, University of Heidelberg, Germany; Department of Infectious Diseases, Molecular Virology, University of Heidelberg, Germany; Department of Molecular Genetics, Weizmann Institute of Science, Rehovot, Israel; Department of Molecular Biology, University Medical Center Göttingen, Göttingen, Germany; Medical Biochemistry and Molecular Biology, Center for Molecular Signaling (PZMS), Saarland University, Homburg/Saar, Germany; Departments of Dermatology, Venereology, Allergology and Immunology, Dessau Medical Center, Brandenburg Medical School Theodor Fontane Dessau, Germany; Medical Institute of Bioregulation, Kyushu University, Fukuoka, Japan

## Abstract

Fatty acyl-CoA reductase 1 (Far1) is an ubiquitously expressed peroxisomal membrane protein generating fatty alcohols required for the biosynthesis of ether lipids.

Lipid droplet localization of human Far1 was observed by fluorescence microscopy under conditions of increased triglyceride synthesis in tissue culture cells. This unexpected finding was supported further by correlative light electron microscopy and subcellular fractionation. Selective permeabilization and N-glycosylation tagging suggest that Far1 is able to assume two different membrane topologies, differing in the orientation of the short hydrophilic C-terminus towards the lumen or the cytosol, respectively. Two closely spaced hydrophobic domains are contained within the C-terminal region. When analyzed separately, the second domain was sufficient for the localization of a fluorescent reporter to lipid droplets. Targeting of Far1 to lipid droplets was not impaired in either pex19 or TRC40/ASNA1 CRISPR/Cas9 knockout cells.

In conclusion, our data suggest that Far1 is a novel member of the rather exclusive group of dual topology membrane proteins. At the same time, Far1 shows lipid metabolism-dependent differential subcellular localizations to peroxisomes and lipid droplets.

## Introduction

Fatty acids are a fundamental requirement for the biogenesis of membranes, and their oxidation generates efficiently energy in the form of ATP. It is therefore not surprising that almost all cells have the capacity to store fatty acid derivatives in specialized organelles now widely termed lipid droplets (LDs; Murphy, 2012; Walther and Farese Jr, 2012). LDs are highly dynamic organelles, with their size and number dependent on nutrient supply and hormonal cues. Neutral lipids, such as triglycerides and cholesteryl esters, are stored in the hydrophobic core, which is surrounded by a phospholipid monolayer with embedded proteins. LD proteomics of mammalian cells revealed the presence of constitutively associated LD proteins (perilipins) and many enzymes of lipid metabolism (Bersuker et al., 2018; Brasaemle et al., 2004; Goodman, 2009; Liu et al., 2004). Hydrophobic domains are somehow required for targeting of proteins to lipid droplets, but the mechanisms involved appear to be diverse and not fully understood (Kory et al., 2016; Ohsaki et al., 2014; Thiele and Spandl, 2008).

Fatty acyl-CoA reductases (Far) are peroxisomal membrane proteins that catalyze the formation of fatty alcohols from fatty acyl-CoA and NADPH (Cheng and Russell, 2004), which in turn are utilized for ether lipid synthesis. In mammals, the Far family is represented by Far1 and Far2, which show a high overall homology of 58% (Cheng and Russell, 2004). Far1 expression has been found in most of the tissues whereas the distribution of Far2 mRNA is more restricted (Cheng and Russell, 2004). Patients with a homozygous knockout of the FAR1 gene suffer from severe epilepsy, cataracts and profound growth and mental retardation (Buchert et al., 2014), suggesting that Far2 cannot compensate for Far1 at the whole organism level. Most peroxisomal membrane proteins are synthesized on free ribosomes in the cytosol and are inserted into the peroxisomal membrane either by direct targeting (class I proteins) or via the ER (class II proteins) (Smith and Aitchison, 2013).

Membrane protein topology describes the number and orientation of one or more hydrophobic domains relative to the membrane (von Heijne, 2006). The vast majority of membrane-associated proteins possesses only one stable configuration. However, there is a small subset of proteins that show changes in the orientation of one, multiple or all transmembrane helices upon membrane insertion, resulting in complex topological variants (von Heijne, 2006). The phenomenon of multiple topologies of a single protein is very rare and neither the evolutionary background nor the mechanisms of exhibiting multiple topological variants are understood satisfyingly. Here, we demonstrate that Far1 is also located on lipid droplets, and suggest that this is due to the presence of two different topological orientations.

## Materials and Methods

### Cell culture, transient and stable expression

Human osteosarcoma U-2 OS cells were cultured in McCoys 5A modified medium (Thermo Fisher Scientific, Waltham, MA, USA) supplemented with 10% FCS (Life Technologies, Carlsbad, CA, USA), 2 mM L-Glutamine (GlutaMAX, Life Technologies, Carlsbad, CA, USA) and 1% penicillin/streptomycin (Life Technologies, Carlsbad, CA, USA). Cells were subcultured when reaching 80% confluence. Human immortalized SZ95 sebocytes (Zouboulis et al., 1999) were cultured in Sebomed^®^ Basal medium (Merck, Kenilworth, NJ, USA) supplemented with 10% FCS, 1 mM CaCl_2_ and 5 ng/ml hEGF.

For transient transfection cells were grown on coverslips to 80-90% confluence and were transfected with FugeneHD (Promega, Madison, WI, USA) according to the manufacturer’s protocol. Briefly, the transfection mix for a 12 well plate with 0.8 ml medium included 40 μl of OptiMEM (Life Technologies, Carlsbad, CA, USA), 0.8 μg total DNA and 4 μl FugeneHD (Promega, Madison, WI, USA). For 6-well plates, the ratio was adjusted to 2 ml medium. After 4 h, the transfection mix was removed and cells were cultured overnight in antibiotic-free medium. Lipid droplet growth was induced by adding 600 μM oleate (from a 8.66 mM OA:BSA stock solution; molar ratio 6:1; oleic acid from Sigma O1383, BSA from Sigma A8806, St. Louis, MO, USA). The OA-medium was present on the cells overnight, if not indicated otherwise.

For stable expression, the constructs were integrated into the genome by retroviral integration as described elsewhere (Schuck et al., 2004). Briefly, phoenix-gp cells were transfected with the indicated fusion proteins cloned into the retroviral pRIJ vector (Küch et al., 2014) or pRIJ-neo vector. Viral particles contained in the media supernatants were harvested and applied to the cells, followed by antibiotic selection of the transduced target cells (puromycin: 2 μg/ml, 48 h; neomycin: 2 mg/ml for 4 days). Cells were never allowed to reach densities above 90% during the selection process.

### Plasmids

Full length Far1 cDNA was obtained from a cDNA library of SZ95 sebocytes (Zouboulis et al., 1999) and was cloned with PacI and KpnI into a modified version of pEGFP-N1 (Clontech) containing a FLAG epitope at the 3´-end of eGFP (Far1-GFP-FLAG). Sequencing confirmed the cDNA to be 100% identical to the database entry (NM_032228.5).

For the N-terminally tagged version, the full length coding region of Far1 was amplified, cloned it into mCherry-C1 (Clontech) and subsequently subcloned yielding FLAG-mCherry-Far1.

FLAG-mCherry-Far1 was used as a template for all further cloned deletion mutants (Fig. 3B), double epitope tag variants (Fig. 2 and Fig. 4) and the opsin tagged variants (Fig. 2).

The N-terminus of ACSL3 (aa 1-135 of human ACSL3) fused to GFP was used as a lipid droplet marker protein (Poppelreuther et al., 2012). As an endoplasmic reticulum stain, a GFP endowed with the CD8 signal sequence and a KDEL ER-retaining sequence was transfected. Alternatively, the endoplasmic reticulum resident acyl-CoA-synthetase FATP4 (Milger et al., 2006) was used. Fluorescent proteins containing a C-terminal SKL tag (GFP-SKL and BFP-SKL) were used as lumenal peroxisomal marker proteins. The full length cDNA of human pex11β fused to GFP or mCherry, respectively, was used as a marker for peroxisomal membranes (Koch et al., 2010).

More details on the plasmid sequences, cloning strategies and primers are freely available upon request. All constructs created were sequenced for validation.

### Immunofluorescence and microscopy

U-2 OS cells were grown on 10 mm coverslips and were fixed with 4% paraformaldehyde in PBS for 20 minutes. Cells that were not processed for immunofluorescence were directly embedded in MOWIOL 4-88 (Calbiochem, San Diego, CA, USA).

For conventional immunofluorescence, the cells were permeabilized with 0.1% saponin and blocked with 0.5% gelatin and 0.5% BSA (SGB) for 10 min at room temperature. The cells were incubated with the first antibody diluted in SGB for 1h. The coverslip was subsequently washed 3x with SG-buffer (0.1% saponin, 0.5% gelatin) and the secondary antibodies were diluted in SGB and applied for 60min. After two washes with SG-buffer and two washes in PBS the cells were embedded in MOWIOL.

For immunofluorescence under selective permeabilization conditions, cells were fixed as described above and permeabilized with 10 μM digitonin for 10 min on ice. Permeabilization with 1% Triton-X 100 on ice solubilized all membranes and served as a staining control. Cells were subsequently blocked with 1% BSA for 1 h at room temperature and incubated with the first antibody diluted in 1% BSA for 1 h. Cells were then washed three times for five minutes with PBS and incubated with the secondary antibody diluted in PBS for 1 h at room temperature. After additional three washes for five minutes in PBS, the cells were embedded in MOWIOL. The intraluminal ER protein CaBP1 (Fullekrug et al., 1994) was used to verify the inaccessibility of membrane covered epitopes after digitonin permeabilization.

Neutral lipids were stained with 2 μg/ml BODIPY 493/503 (Stock 1 mg/ml in EtOH; Invitrogen D-3922, Waltham, MA, USA) or 1 μM MDH (Abgent, SM1000a, San Diego, CA, USA; Stock 1 mM) diluted in PBS for 15 min at room temperature.

Raw images were acquired on an Olympus BX41 microscope, 60x oil immersion Plan S Apo NA 1.35, F-view II CCD camera, cell^D software.

Pictures were taken adjusting brightness and gamma. Gamma values reached from 1.5 (ER structures) over 1.3 (peroxisomes) to 1.0 (lipid droplets).

Adobe Photoshop was used to arrange the figure, colorizing grey value pictures and adjust brightness.

### Antibodies

Immunofluorescence was performed with rabbit anti-CaBP1 (Fullekrug et al., 1994), rabbit anti-FLAG (Sigma F7425, St. Louis, MO, USA; 1:1000 for IF) and mouse anti-HA (Santa Cruz SC-7392, Dallas, TX, USA; 1:200 for IF). As a lipid droplet marker, anti-TIP47 antibody (R&D Systems, Minneapolis, MN, USA), 1:600) was used.

For western blot, mouse anti-FLAG (Sigma F1804 St. Louis, MO, USA; 1:4000), rabbit anti-HA (Santa Cruz SC-805 Dallas, TX, USA; 1:1000) and mouse anti-actin (Sigma A5441 St. Louis, MO, USA; 1:40.000) were used. As markers for the ER, anti-calnexin (Thermo Fisher Scientific, Waltham, MA, USA, 1:200) was detected. Anti-ACSL3 (Abnova, Taipei, Taiwan, 1:3000) served as a marker for lipid droplets.

Anti-pex19 antibody (Thermo Fisher Scientific, Waltham, MA, USA) was used in a 1:10.000 dilution for western blot.

### Correlative light electron microscopy (CLEM)

Cells were seeded at a low density in MatTek dishes (Ashland, MA, USA) having glass gridded coverslips with an alphanumeric pattern for relocating cells or cells clusters. After treatment, cells were fixed with 4% PFA and 0.2% glutaraldehyde in PBS for 30 min. After fixation, the cells were washed three times with 150 mM glycine and twice with PBS. Subsequently cells were imaged with a Nikon Ti Eclipse microscope equipped with an Ultraview VoX confocal spinning disc system (PerkinElmer, Waltham, MA, USA). First, cells were imaged at 20X magnification (20X objective, Plan Fluor, NA 0.75, oil), to record the positions of the cells of interest expressing fluorescent proteins. Then 60X magnification images (60X objective: Apo TIRF, NA 1.49, oil) were also acquired to improve the correlation with the electron microscopy (EM) data. Post-fixation of the cells was carried out with 2% osmium tetroxide (OsO_4_) in 50mM Na-cacodylate buffer for 40 min on ice. Afterwards, the cells were washed three times with ddH_2_O and stained with 0.5% uranyl acetate (UA) in H_2_O for 30 min. After three additional washes with ddH_2_O the cells were dehydrated in an increasing ethanol series at room temperature (40%, 50%, 60%, 70% and 80%) for five minutes each followed by 95% and 100% ethanol for 20 min each. The cells were then quickly immersed in 100% propylene oxide and embedded in an Araldite-Epon mixture (Araldite 502/Embed 812 kit, Electron Microscopy Sciences, Hatfield, PA, USA). The samples were incubated for 48 h at 60°C for polymerization of the resin. The MatTek dishes and the gridded coverslips were subsequently removed from the polymerized resin blocks and the embedded cell monolayers were sectioned using a Leica Ultracut UCT ultramicrotome (Leica Microsystems) and a diamond knife (Ultra 35 ° from Diatome). Finally the 70 nm sections were counter-stained with 2% UA in 70 % methanol and lead citrate in CO_2_-free H_2_O and visualized under a JEM-1400 (JEOL) TEM equipped with a TemCam-F416 digital camera system (TVIPS) operated at 120 kV. Correlations between the light microscopy images and the electron micrographs were calculated using the non-rigid transformation of the Icy ec-CLEM plugin (http://icy.bioimageanalysis.org/) (Paul-Gilloteaux et al., 2017). CLEM as applied here has been described in detail recently (Romero-Brey, 2018).

### Subcellular fractionation

Lipid droplets were prepared by a sucrose step gradient (modified after (Thul et al., 2017) and (Krahmer et al., 2013)). For subcellular fractionation, cells were grown subconfluent in two 145 cm^2^ plates and were treated with or without 600 μM oleic acid overnight. The following steps were carried out at 4°C. Cells were washed three times with PBS, collected by scraping and pelleted. The pellet was resuspended in buffer A1 (50 mM TRIS (pH 7.4), 20 mM sucrose, 1 mM EDTA, 1 mM β-mercaptoethanol in H_2_O) and homogenized using 10 strikes with a G22 needle and additional 23 strikes with a G25 needle. Centrifugation at 1000 g for 10 min obtained the postnuclear supernatant (PNS). The PNS was centrifuged at 100.000xg (TLA55 rotor) for 1 h to obtain a membrane pellet and the cytosolic supernatant including lipid droplets. To further separate the cytosol from lipid droplets, the supernatant was adjusted to 2.7 ml with buffer A1 and mixed with 2.7ml of buffer A4 (50 mM Tris pH 7.4, 1.08 M Sucrose, 1 mM EDTA, 1 mM β-mercaptoethanol in H_2_O). Together, it formed the bottom layer of the sucrose gradient. This was carefully overlayed with 1.8 ml of buffer A3 (50 mM Tris pH 7.4, 270 mM Sucrose, 1 mM EDTA, 1 mM β-mercaptoethanol in H_2_O), buffer A2 (50 mM Tris pH 7.4, 135 mM Sucrose, 1 mM EDTA, 1 mM β-mercaptoethanol in H_2_O) and buffer A1. This step gradient was centrifuged for 1:45 h at 150.000xg in an SW41TI rotor. Fractions were collected from top to bottom in 1.35 ml steps. In the end, 8 fractions were obtained from the gradient. Proteins of fraction 1-8 were precipitated with chloroform: methanol 1:2 (v/v) and resuspended in 40 μl sample buffer (2% SDS (w/v), 62.5 mM Tris pH 6.8, 10% (v/v) glycerol, 100 mM β-mercaptoethanol). The membrane pellet was directly dissolved in 320 μl sample buffer. 10 μl of each fraction were subjected to SDS-PAGE with subsequent western blot analysis. After oleic acid incubation the enrichment compared to the ER marker protein calnexin was 17.2 for ACSL3, 12.4 for Far1 and 4.1 for pex11β.

### Yeast strains and transformations

For evaluation of hsFar1 localization in yeast, the cDNA coding for FLAG-mCherry-Far1 was cloned into pYX142 (R&D Systems, Minneapolis, MN, USA) using PCR primers encoding XmaI and NheI restriction sites (s_BglII_XmaI_FLAG *5’-acgtagatctcccggg-accatggactacaaggacgacg-3’* and a_NheI_XbaI_NotI_Far1Ct *5’-acgtgctagctctagagcggccg-ctcagtatctcatagtgctgg-3’*). For the microscopy analysis, the lipid droplet marker Erg6 (Pu et al., 2011) and the peroxisomal marker Pxa1 (Shani and Valle, 1996) in the yeast strains BY4741 GFP-Pxa1 (his3∆1 leu2∆0 met15∆0 ura3∆0 hph∆n::URA3::SpNOP1pr-GFP-Pxa1) and BY4741 GFP-Erg6 (his3∆1 leu2∆0 met15∆0 ura3∆0 hph∆n::URA3::SpNOP1pr-GFP-Erg6) were used. Liquid cultures were grown in overnight at 30°C in SD-URA medium. Strains were diluted to an OD600 of ~0.2 into a new plate containing SD medium (6.7 g/L yeast nitrogen base and 2% glucose) or S-Oleate (6.7 g/L yeast nitrogen base, 0.2% oleic acid and 0.1% Tween-80). Strains were incubated at 30°C for 4 h in SD medium or for 20 h in S-Oleate. The cultures in the plates were then transferred into glass-bottom 384-well microscope plates (Brooks life science systems, Chelmsford, MA, USA) coated with concanavalin A (Sigma, St. Louis, MO, USA). After 20 min, wells were washed twice with SD-riboflavin complete medium (for strains in glucose) or with double-distilled water (for strains in oleate) to remove non-adherent cells and to obtain a cell monolayer. The plates were then transferred to the ScanR inverted fluorescent microscope system (Olympus, Tokyo, Japan). Images of cells in the 384-well plates were recorded in the same liquid as the washing step at 24°C using a 60x air lens (NA 0.9) and with an ORCA-ER charge-coupled device camera (Hamamatsu, Hamamatsu, Japan). Images were acquired in two channels: GFP (excitation filter 490/20 nm, emission filter 535/50 nm) and mCherry (excitation filter 572/35 nm, emission filter 632/60 nm). All images were taken at a single focal plane.

### Pex19-knockout cell line

For generating the pex19 knockout cell lines, a modified version of the eSpCas9(1.1) ((Slaymaker et al., 2016) Addgene # 71814) equipped with a puromycin resistance gene was used. The guide for pex19 (Schrul and Kopito, 2016) was generated by annealing sense (*5´- caccgtgtcggggccgaagcggac-3´)* and antisense (*5´- aaacgtccgcttcggccccgacac-3´*) oligo and ligation into the backbone containing the Cas9 cut with BbsI. U-2 OS cells were seeded in 6 well plates and transfected with the Cas9 vector equipped with the guide RNA. A Cas9 vector without a guide served as a control. 24 h post transfection the cells were trypsinized and selected with 2 μg/ml puromycin for 48 h. After additional 24 h, cells were trypsinized and 30 cells were seeded in 60 cm^2^ dishes. Two weeks after transfection, single cell clones were transferred to a 24 well plate. Single cell clones were screened by eGFP-SKL transfection for the absence of peroxisomes and finally verified by western blotting.

### ASNA1/TRC40-knockout pool

Knockout of ASNA1 was achieved by transfection of a plasmid containing SpCas9 ((Ran et al., 2013) Addgene #62988) and a guide RNA targeting exon 3 of the TRC40-gene (exon 3 sense *5´- caccgcctgacgagttcttcgagg-3´*antisense *5´-aaaccctcgaagaactcgtcaggc-3´*). After puromycin selection (2 μg/ml) for 48 h the cells were grown for two weeks with cell densities above 30%. No formation of single cell clones could be achieved. For the preparation of genomic DNA, 50.000 cells were resuspended in 17 μl ddH_2_O and 2 μl of 10x Taq-PCR-Buffer (QIAGEN, Hilden, Germany) as well as 800 U proteinase K (New England Biolabs, Ipswich, MA, USA). Incubation at 65°C for 60 min was followed by 15min at 95°C for inactivation of the proteinase K. The targeted genomic region was amplified with conventional PCR (QIAGEN, Hilden, Germany) and sequenced. To estimate the knockout efficiency of the guide in the pool the TIDE algorithm was used (Brinkman et al., 2014) comparing the chromatograms derived from the wildtype locus and the guide-targeted locus. The knockout was further confirmed by western blot and was estimated effective for over 95% of the cells. Antibodies used were rabbit anti TRC40 (Proteintech 15450-1-AP), and rabbit anti tubulin (Proteintech 11224- 1-AP). The antibody incubation was carried out in PBS-0.1% Tween-20 (overnight at 4°C). Blots were imaged using an Odyssey Sa Infrared imaging system with IRDye LiCOR secondary antibodies.

### Software

Guides for ASNA1 were designed with CRISPOR (http://crispor.tefor.net/). Genomic alterations were estimated with the TIDE algorithm (Brinkman et al., 2014). Hydrophobicity scores were generated and evaluated by TMHMM 2.0 (http://www.cbs.dtu.dk/services/TMHMM/); (Krogh et al., 2001). The helical wheel projection was generated by HeliQuest (http://heliquest.ipmc.cnrs.fr/). Adobe Illustrator was used for the schematic models and Adobe Photoshop was used to arrange the figures.

## Results

The biochemistry of long chain fatty acids is at the heart of the complex lipid metabolic network. Here, we analyzed the localization and targeting of fatty acyl-CoA reductase 1 (Far1) in human osteosarcoma U-2 OS cells to gain further insights into the topology and subcellular compartmentalization of fatty acid metabolism.

### Localization of Far1 to peroxisomes and lipid droplets

Fatty acyl-CoA reductases are not widely studied, and there were no suitable antibodies for immunocytochemistry protocols available. Therefore, we cloned the cDNA of Far1 from the human sebaceous gland cell line SZ95 and designed a mCherry-Far1 fusion plasmid. Transient expression in U-2 OS cells showed numerous dot-like structures throughout the cytoplasm which overlapped with the peroxisomal membrane marker GFP-pex11β (Fig. 1A), consistent with earlier work (Honsho et al., 2013).

**Figure 1:**
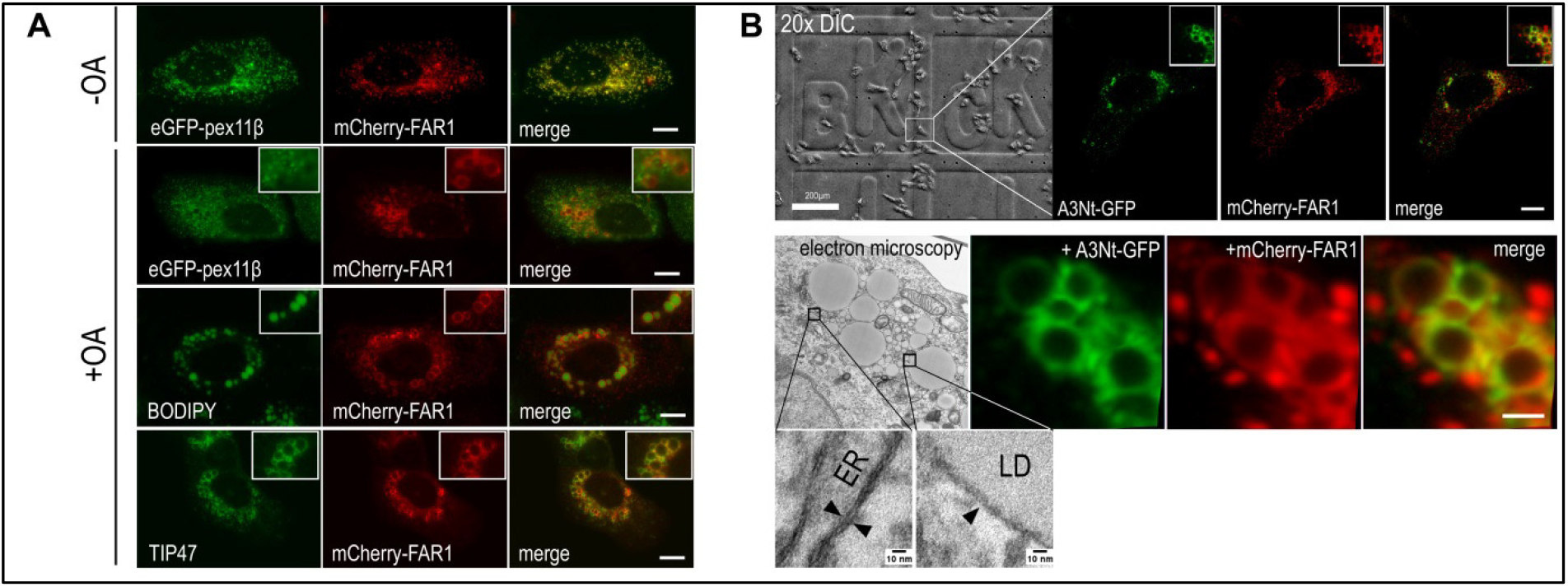
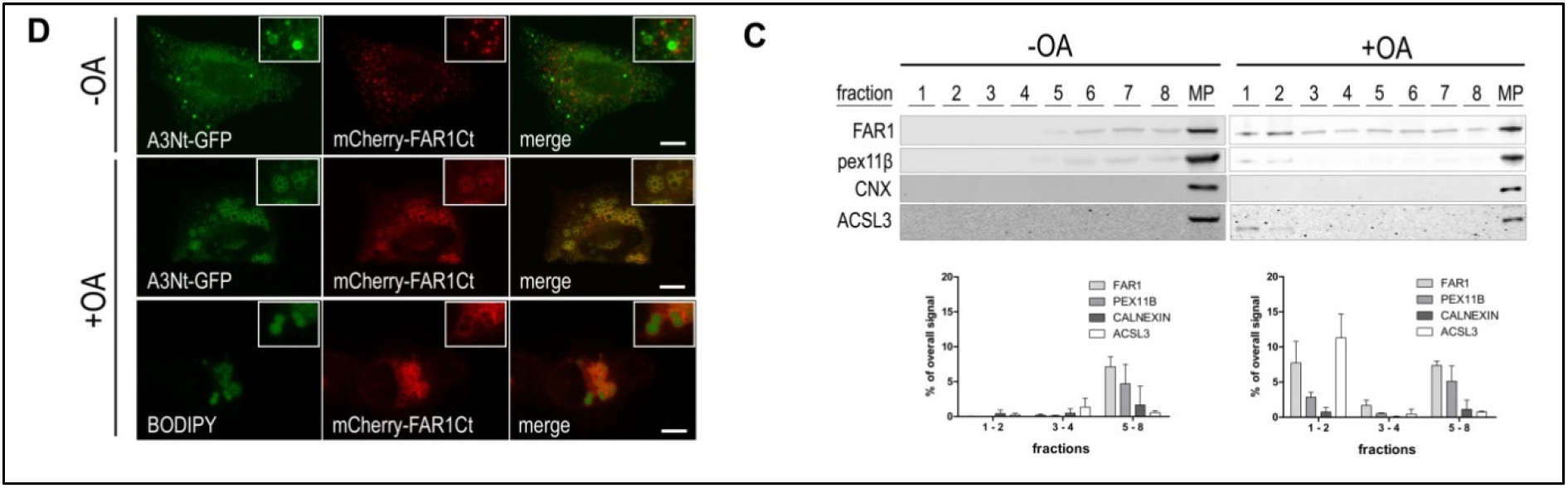
Localization of Far1 to lipid droplets. (A) Localization of Far1 to lipid droplets in fatty acid treated human U-2 OS cells by fluorescence microscopy. Transient expression of mCherry-Far1 (red) and comparison to the peroxisomal membrane marker protein GFP-pex11β (green), the lipid droplet marker Plin3/TIP47 (antibody staining; green, Alexa488) and the neutral lipid stain Bodipy 493/503 (green). Over-lap of mCherry-Far1 and GFP-pex11β indicates peroxisomal localization of Far1 (first panel) using standard growth medium (-OA). After oleic acid incubation (+OA) Far1 segregates from pex11β (second row) and overlaps with endogenous Plin3/TIP47 instead (fourth row); Far1 surrounds lipid droplets stained by the neutral lipid dye Bodipy 493/503 (BODIPY; third row). Scale bars 10 μm. (B) Correlative light electron microscopy (CLEM). U-2 OS cells stably co-expressing mCher-ry-Far1 (red) and the lipid droplet marker A3Nt-GFP (green) were seeded onto glass-gridded MatTeK dishes and supplemented with oleate o/n. DIC (differential interference contrast) was used to determine the cell position on the MatTek dish grid (first row, left). LD localization of Far1 is suggested by the colocalization with A3Nt-GFP by confocal microscopy (first row, right; scale bar 10 μm). Analysis of the same cell by transmission electron microscopy of ultrathin resin sections shows lipid droplets corresponding to the A3Nt-GFP and mCherry-Far1 images by confocal microscopy (second row; bar 1 μm). Higher magnification electron micrographs demonstrate that the resolution is sufficient to identify ER tubules with bilayer membranes and LD monolayers (arrowheads; third row). (C) Subcellular fractionation. U-2 OS cells stably expressing 3xFLAG-GFP-Far1 and 3xHA-mCherry-pex11β were treated with oleate-BSA o/n (+OA) or not (-OA). Heavy membranes were removed first by 100,000xg for 1 h (membrane pellet, MP). Sucrose density gradient centrifugation of the LD enriched supernatant allowed further separation into eight fractions collected from top to bottom (1-8). The LD association of Far1 was assessed by western blotting in comparison to pex11ß, endogenous ACSL3 (LD marker) and calnexin (CNX; ER contaminant). Far1 was enriched 12-fold over calnexin in the LD fractions. Error bars are the SD from n=3 independent experiments. (D) The C-terminus of Far1 is sufficient for lipid droplet localization. The mCherry-Far1Ct fusion protein (Far 1 amino acids 451-515) is distinct from the LD marker A3Nt-GFP under standard conditions (-OA), but overlaps extensively after oleate supplementation (+OA). Neutral lipid staining (BODIPY) is surrounded by mCherry-Far1Ct (third row). Bar, 10 μm.

Far1 localized to lipid droplets after oleate (OA) supplementation, as indicated by the colocalization with the lipid droplet marker Plin-3/TIP47, and the segregation from pex11ß (Fig. 1A). This surprising finding was further confirmed by neutral lipid staining demonstrating an abundance of lipid droplets surrounded by mCherry-Far1 (Fig. 1A).

To investigate this in more detail, we established U-2 OS cells stably expressing mCherry-Far1 together with the lipid droplet marker A3_Nt_-GFP (Poppelreuther et al., 2012) by retroviral transduction. Analysis by correlative light electron microscopy (CLEM) showed that the fluorescent signal of both mCherry-Far1 and A3Nt-GFP corresponded to lipid droplets, which were identified by transmission electron microscopy (TEM) as large circular homogenously electron dense structures surrounded by a more electron dense line (Fig 1B). ER tubules were identified by EM but did not correspond to the circular fluorescence pattern of Far1 and A3Nt-GFP. Additionally, mCherry-positive structures were also associated with electron dense stratified organelles presumably corresponding to peroxisomes.

To enable the direct comparison of two peroxisomal membrane associated proteins by sub-cellular fractionation, epitope tagged variants of Far1 and pex11ß were stably co-expressed by retroviral transduction (3xFLAG-GFP-Far1.3xHA-mCherry-pex11β.U-2 OS cells). Lipid droplets were enriched by sedimenting most of the other cellular membranes for 1 h at 100,000xg. The supernatant was further subfractionated by sucrose density gradient centrifugation and analyzed by western blotting (Fig. 1C). Far1 was clearly enriched in the LD fractions compared to pex11ß, even if it did not reach the levels of our lipid droplet marker protein (endogenous ACSL3).

The 65 C-terminal amino acids of Far1 were sufficient to allow targeting of a fluorescent reporter to lipid droplets in oleate treated U-2 OS cells (mCherry-Far1Ct; Fig. 1D). The same C-terminal region is also necessary for the targeting of Far1 to peroxisomes ((Honsho et al., 2013), and not shown).

### Two different topologies of Far1

On peroxisomes, Far1 displays a classical type I membrane topology with most of the protein including the enzyme domain exposed to the cytosol, and a short luminal tail (Honsho et al., 2013). This topology however is not found on lipid droplet associated proteins because the hydrophobic core does not allow penetration by charged amino acids (Kory et al., 2016). In addition, lipid droplets are surrounded by a phospholipid monolayer which is too narrow to incorporate standard integral transmembrane bilayer domains.

To resolve this apparent conflict, we used double epitope tagging of Far1 (FLAG-Far1-HA) combined with selective permeabilization. Digitonin solubilizes the plasma membrane but not intracellular membranes like the ER or peroxisomes, allowing access of antibodies only to cytoplasmically oriented epitopes. Antibodies against the C-terminal HA epitope tag stained Far1 only when it was present on lipid droplets in oleate treated cells, or if the non-discriminative detergent Tx-100 was used (Fig. 2A). This suggested that the orientation of the C-terminus would be towards the cytosol on lipid droplets but extracytosolic/intralumenal on peroxisomes.

**Figure 2:**
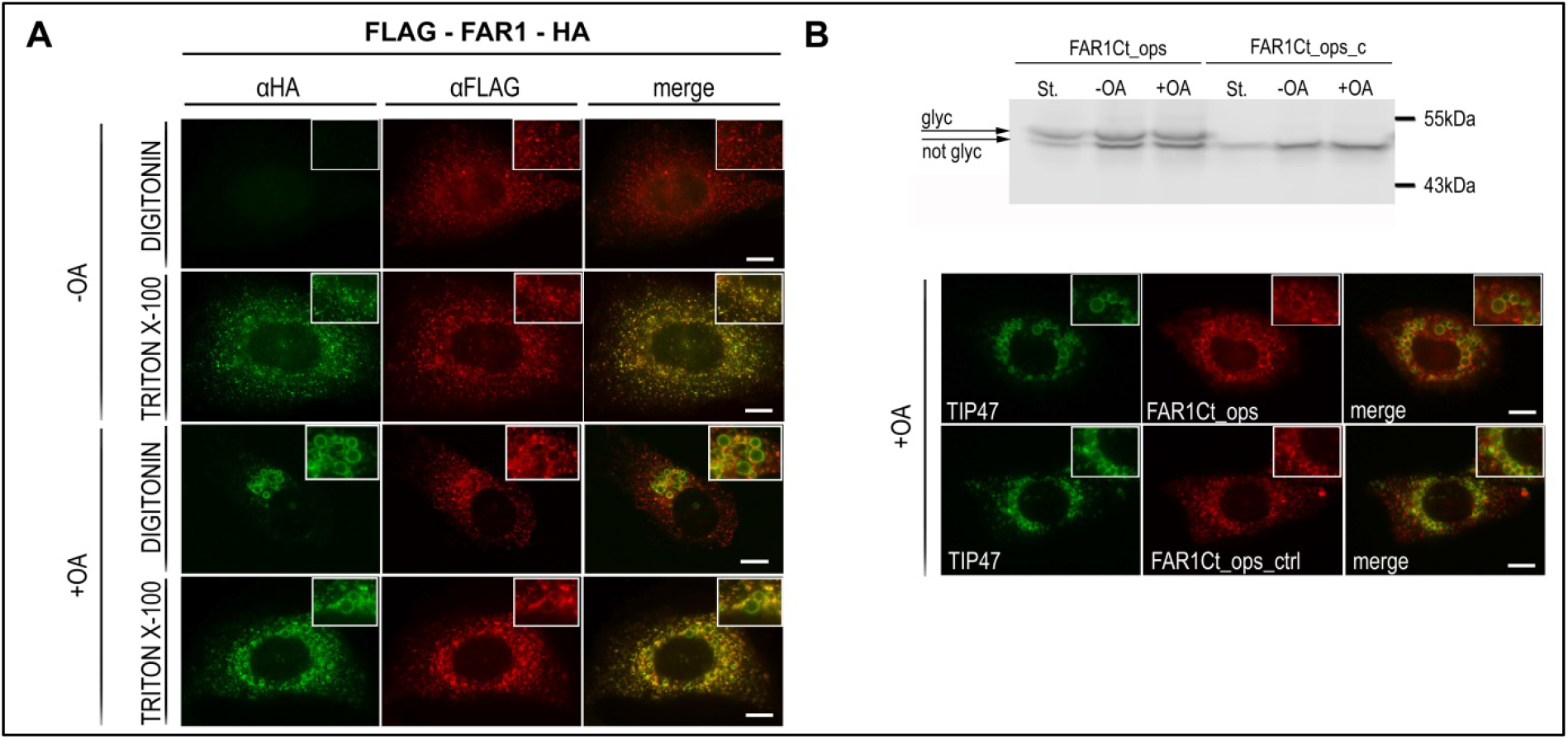
Localization dependent topology of Far1. (A) Selective permeabilization of U-2 OS cells expressing double epitope tagged Far1. The Far1 construct features an N-terminal FLAG and a C-terminal HA epitope tag; see also Fig. 3B. Digitonin permeabilizes the plasma membrane but no intracellular membranes, allowing access of antibodies only to cytoplasmatically oriented epitopes. Indirect immunofluorescence with anti-FLAG (red) and anti-HA (green) antibodies indicated that the C-terminus of Far1 is oriented towards the inside of the peroxisomes (first row) at standard growth conditions (-OA). After oleate supplementation (+OA), HA antibodies efficiently stained lipid droplets (third row), suggesting that both N- and C-terminus are exposed to the cytosol. Note that remaining peroxisomal Far1 was not stained by the HA antibodies (third row). Triton-X-100 solubilizes all membranes indiscriminately and was used as a staining control (second and fourth row). Bar 10μm. (B) N-glycosylation tagging of the Far1 C-terminus. The opsin oligopeptide tag (ops) contains two consensus sites for N-glycosylation whereas the corresponding control peptide (ops_ctrl) cannot be glycosylated. Western blotting and detection by anti-FLAG indicated partial glycosylation under all conditions (St. starved cells without any lipid droplets; -OA standard growth medium; +OA supplementation with oleate o/n). Both Far1Ct_ops and Far1Ct_ops_ctrl (red, anti-FLAG) localized partially to lipid droplets as indicated by the overlap with endogenous Plin3/TIP47 (green). Bar 10μm.

Targeting of LD proteins starts either from the ER or from the cytosol (Kory et al., 2016). To distinguish between these possibilities, we applied glycosylation tagging of Far1 with an opsin derived sequence (Bano-Polo et al., 2011; Favaloro et al., 2008; Nilsson and von Heijne, 1993). Stably expressing cells were analyzed by western blotting and immunofluorescence (Fig. 2B). Partial glycosylation of Far1 constructs was observed only with intact N-glycosylation motifs, as compared to the opsin control sequence containing two point mutations. Remarkably, glycosylation was not only present in oleate treated cells but also if cells underwent starvation to remove lipid droplets. This suggested that in our model system, the ER would also be a part of the trafficking itinerary towards peroxisomes. Both opsin constructs were partially found on lipid droplets (Fig. 2B), likely corresponding to the non-glycosylated variants because N-glycoproteins are neither expected to be found on LDs nor has there been any report of this to the best of our knowledge.

### The C-terminus of Far1 contains two distinct membrane association domains

*In silico* transmembrane domain prediction by TMHMM (Krogh et al., 2001) suggested two C-terminal hydrophobic domains (HDs), of which the first one (HD1: aa 466-483) reached the threshold for a bona fide transmembrane domain, whereas the second hydrophobic domain (HD2: aa 492-510) did not (Fig. 3A). Starting with mCherry-Far1Ct (Fig. 1D), either HD1 or HD2 were deleted (Fig. 3B) and analyzed for their localization. Both mCherry-HD1 and mCherry-HD2 localized to the ER under steady state conditions (Figs. 3C, 3D). Oleate treatment did not change the localization of mCherry-HD1 but mCherry-HD2 overlapped strikingly with the lipid droplet marker A3Nt-GFP (Fig. 3D). Neither of the constructs localized to peroxisomes, likely because the putative binding site for the peroxisomal sorting chaperone pex19 (Honsho et al., 2013; Van Ael and Fransen, 2006) was compromised.

**Figure 3:**
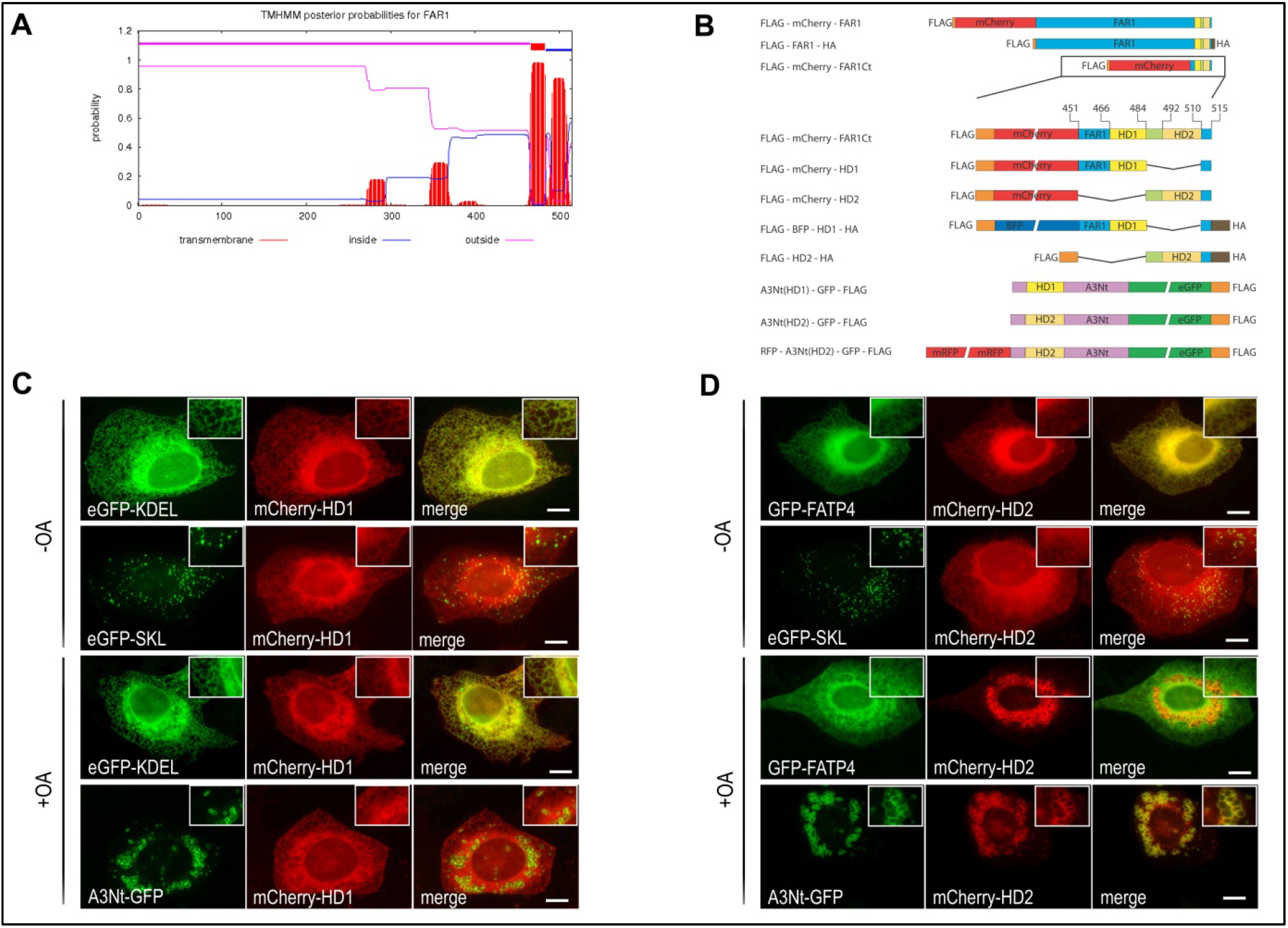
The C-terminus of Far1 contains two membrane association domains. (A) Transmembrane domain prediction of human Far1 (Uniprot Q8WVX9) by the TMHMM Server. Only the first hydrophobic stretch of amino acids (466-484) exceeds the threshold for a bona fide transmembrane domain (red bar). (B) Overview of Far1 constructs. The Far1 C-terminus contains the two hydrophobic domains including the surrounding amino acids (S451 - Y515). HD1 (yellow, I466-A483): first hydrophobic domain; HD2 (light brown, I492-S510): second hydrophobic domain; intermittent amino acids (light green, A484-N491); C-terminal end (light blue, S511-Y515). A3Nt (light purple) corresponds to the N-terminus of ACSL3 (Poppelreuther et al., 2012). (C) The first hydrophobic domain of Far1Ct (mCherry-HD1) confers localization to the endoplasmic reticulum, as indicated by the overlap with the luminal ER marker GFP-KDEL (second and fourth row). It is segregated from both peroxisomes (second row) and lipid droplets (fourth row). (D) The second hydrophobic domain is sufficient for targeting the reporter (mCherry-HD2) to lipid droplets (fourth row). Under standard growth conditions, mCherryHD2 is predominantly localized to the ER but not to peroxisomes, as indicated by the overlap with the ER membrane marker GFP-FATP4 and the segregation from peroxisomal eGFP-SKL(first and second row). Standard growth medium (-OA); oleate-BSA o/n (+OA). Scale bar 10μm.

Both constructs were double epitope tagged (Fig. 3A) and retrovirally transduced into U-2 OS cells, followed by selective permeabilization analysis, as above. The HD1 construct behaved like a type I integral membrane protein of the ER, with the C-terminal HA epitope oriented into the lumen and only accessible using TX-100 to completely permeabilize all subcellular membranes (Fig. 4A). Remarkably, the HD2 construct was stained efficiently when using digitonin, suggesting that both the N- and the C-terminus were facing the cytosol (Fig 4C).

**Figure 4:**
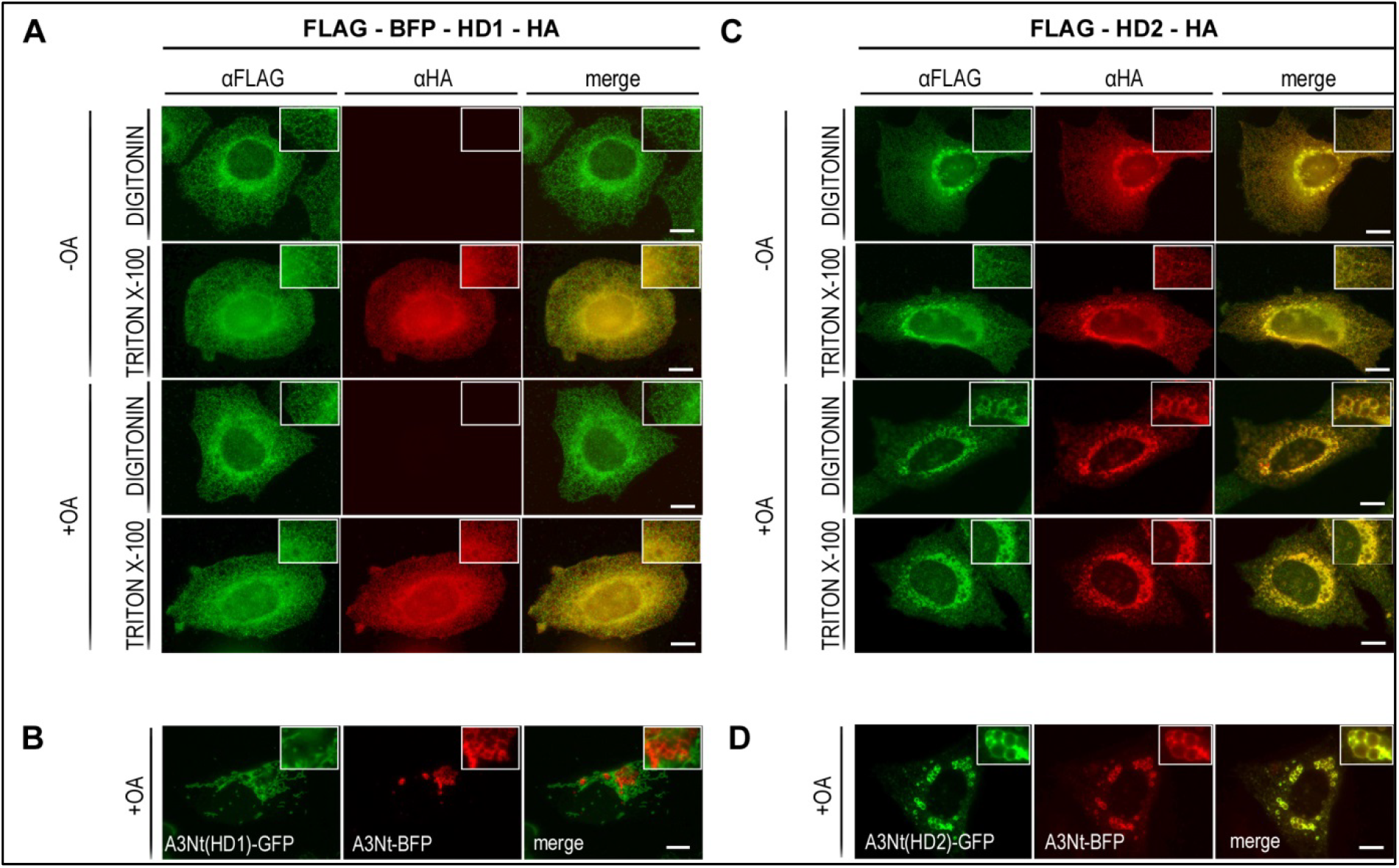
The two hydrophobic domains of Far1 confer different topologies. As before, cells were cultured with standard growth medium (-OA) or supplemented with ole-ate-BSA o/n (+OA). All dimension bars are 10 μm. (A) Selective permeabilization of FLAG-BFP-HD1-HA stably expressed in U-2 OS cells. Permeabilization with digitonin prevented binding of the anti-HA antibodies (αHA) to the corresponding epitope at the C-terminus, suggesting an integral TMD topology. Triton X-100 was again used as a permeabilization control (see also Fig. 2A). (B) A3Nt(HD1)-GFP contains the N-terminal region of ACSL3, but with the hydrophobic domain replaced by the Far1 HD1. This construct localized to mitochondria but not to lipid droplets here marked by A3Nt-BFP; the blue fluorescence was false-colored to red. (C) Selective permeabilization of FLAG-HD2-HA indicates that both N- and C-terminus are oriented towards the cytosol, because permeabilization of the plasma membrane was sufficient for binding of the anti-HA (αHA) antibodies to the C-terminal epitope. FLAG-HD2-HA localized on lipid droplets after oleate supplementation, consistent with Fig. 3D. (D) The hydrophobic domain of ACSL3 was replaced by the Far1 HD2 sequence. The corresponding A3Nt(HD2)-GFP construct localized efficiently to lipid droplets, indistinguishable from the A3Nt-BFP marker.

The HD1 and HD2 mCherry constructs still contained adjacent amino acids from the Far1 sequence. To further analyze the two HDs, only the hydrophobic domains themselves were used to replace the HD of the A3Nt-GFP lipid droplet marker (Fig. 3B). While the hydrophobic domain of ACSL3 allows both the N- and C-terminus to be oriented towards the cytosol at the ER and on LDs (Poppelreuther et al., 2012), it has not been experimentally clarified whether this is due to the formation of an amphipathic helix or a hairpin motif (Kory et al., 2016). Regardless, the corresponding A3Nt-HD2-GFP construct behaved virtually indistinguishable from A3Nt-GFP, overlapping extensively with lipid droplets after oleate incubation (Fig. 4D), and with Sec61ß at the ER under steady state conditions (Fig. S2). A3Nt-HD1-GFP localized to mitochondria, which was consistent with *in silico* predictions for a mitochondrial targeting peptide (http://www.cbs.dtu.dk/services/TargetP/; not shown).

### Targeting of Far1 to lipid droplets is independent of pex19 or ASNA1

The sorting of Far1 to peroxisomes depends on the cytosolic trafficking chaperone pex19 (Honsho et al., 2013). Recently, pex19 has also been implicated in the targeting of UBXD8 towards lipid droplets via the ER (Schrul and Kopito, 2016). To test a possible involvement of pex19 in the targeting of Far1 to lipid droplets, pex19 was inactivated in U-2 OS cells by CRISPR/Cas9 mediated indel formation (Fig. 5A). As expected, peroxisomes were absent as verified here by the cytosolic location of the peroxisomal SKL reporter protein (Fig. 5B). Re-expression of pex19 restored peroxisomal sorting of BFP-SKL (not shown). However, the localization of Far1 on lipid droplets was not impaired (Fig 5B).

**Figure 5:**
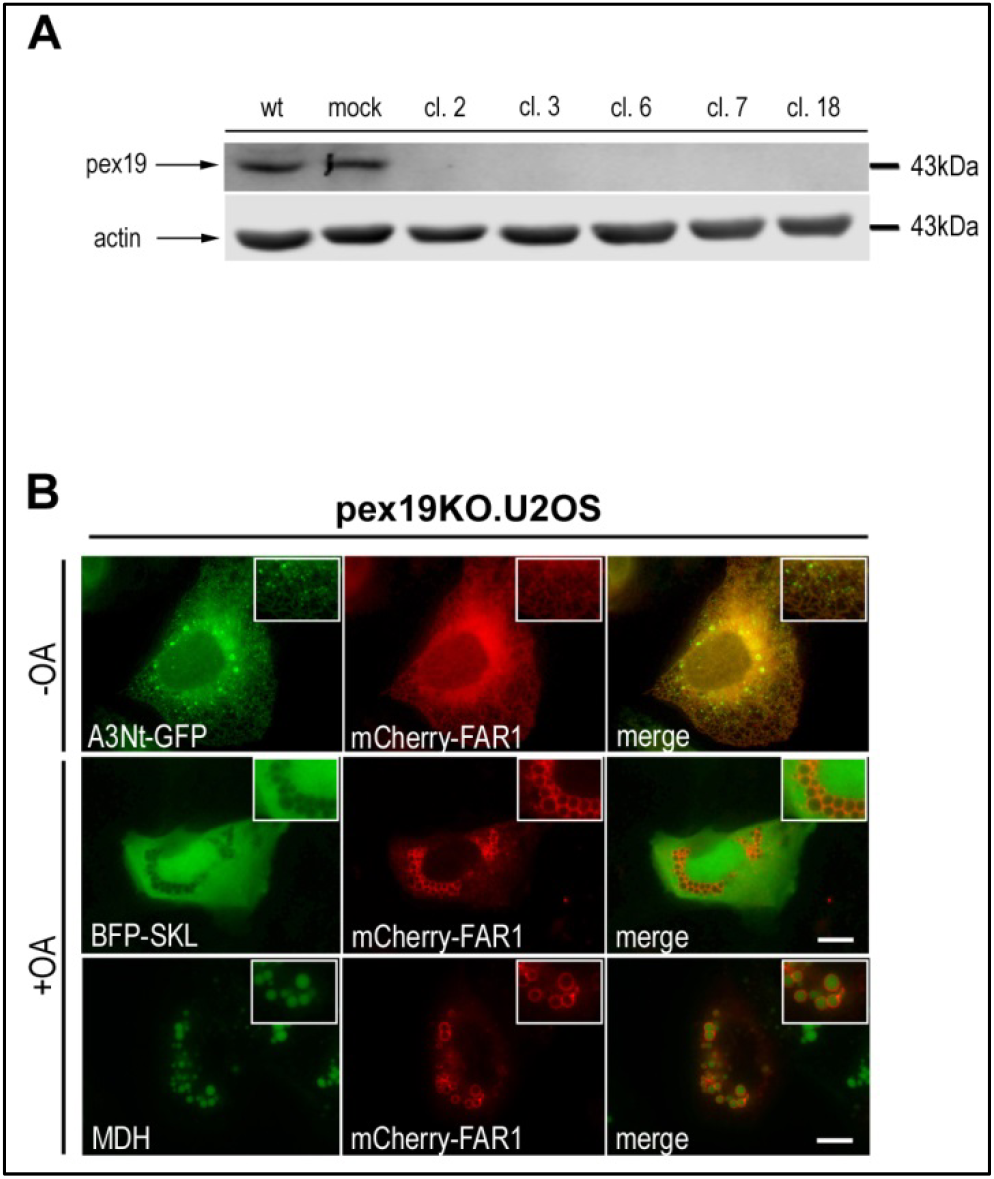
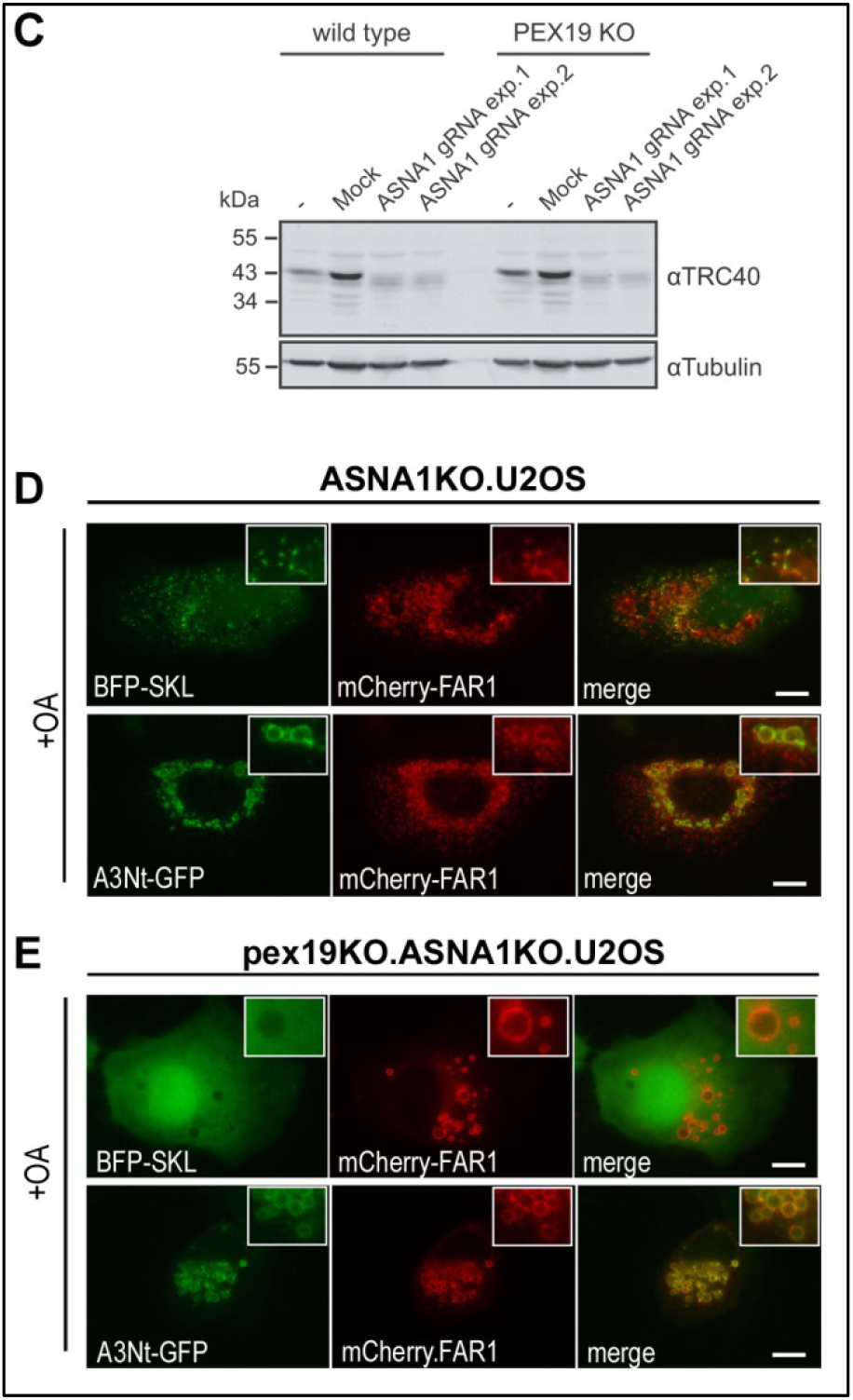
Far1 targeting to lipid droplets is neither dependent on pex19 nor ASNA1. (A) Knockout of pex19 in U-2 OS cells. CRISPR/Cas9 followed by single cell cloning established five cell lines (cl. = clone) deficient for pex19 as shown by western blotting. Mock treated cells were obtained by using the Cas9 plasmid without a guide sequence. ß-actin was used as a loading control. Wildtype cells (wt) are shown as an additional control. (B) Localization of Far1 to the ER and lipid droplets in pex19-KO cells. Far1 (mCherry-Far1, red) surrounding LDs is evidenced by the overlay with the neutral lipid dye MDH (false-colored to green); third row. Without oleate supplementation, Far is retained in the network-like structures of the ER also marked by A3Nt-GFP (first row). Cytosolic fluorescence of the luminal peroxisomal marker BFP-SKL (false-colored to green) indicated the absence of peroxisomes (second row). Standard growth medium (-OA); 600 μM oleate-BSA o/n (+OA). Scale bar 10μm. (C) Knockout of ASNA1 in wt U-2 OS and pex19-KO cells. Antibiotic selection for transient Cas9 expression generated pools of cells which were used here, because no single cell clones could be obtained (see Methods). Western blotting confirmed that the depletion of ASNA1/TRC40 was efficient. Mock: Cas9 plasmid without gRNA transfected; ß-tubulin was used as a loading control. (D) Depletion of ASNA1/TRC40 does not influence targeting of Far1 to peroxisomes or lipid droplets. Far1 overlaps with the peroxisomal marker BFP-SKL (false-colored to green; first row) as well as with LD localized A3Nt-GFP (second row). (E) Localization of Far1 to lipid droplets is not impaired in cells lacking both pex19 and ASNA1. The cytosolic distribution of BFP-SKL confirms the absence of peroxisomes (top row). Far1 overlaps with A3Nt-GFP on lipid droplets in oleate treated (bottom row). (B, D, E) Cells were cultured with standard growth medium (-OA) or supplemented with ole-ate-BSA o/n (+OA). All dimension bars are 10 μm.

ASNA1/TRC40 targets C-terminal anchored membrane proteins to the ER (Borgese and Fasana, 2011; Guna et al., 2018; Rabu et al., 2009). Since the membrane association domains of Far1 are also located at the C-terminus, we considered a role for ASNA1 in the targeting of Far1 (to the ER; further trafficking towards LDs would be mediated either by the HD2 domain or additional factors). For unknown reasons, single cell derived clonal lines could not be obtained after targeting of ASNA1 by CRISPR/Cas9. Instead, cells were selected for the transient presence of Cas9 and propagated as a pool of cells with an estimated overall knockdown efficiency of >95% as assessed by western blotting (Fig. 5D) and genome sequencing (Suppl. Fig. S3). The localization of Far1 on lipid droplets was not changed however (Fig. 5E). The knockdown of ASNA1 was also applied in the pex19-KO background, but again did not change the LD localization of Far1 (Fig. 5F).

## Discussion

### Localization of Far1 to lipid droplets

We demonstrated the presence of Far1 on lipid droplets in U-2 OS cells by several independent methods, and showed that the C-terminal 65 amino acids are sufficient for this unusual localization of a peroxisomal membrane protein (Figs. 1-3). Additionally, Far1 was also observed on LDs in oleate treated epithelial COS-7 cells by fluorescence microscopy, and the closely related Far2 protein was equally localized to LDs (not shown). Consistent with our results, proximity labeling based proteomics suggested that Far1 is the most abundant peroxisomal protein on lipid droplets of U-2 OS cells (Bersuker et al., 2018).

Close *proximity* between peroxisomes and LDs has been observed previously in different model organisms, sometimes enhanced by the remarkable flexibility of peroxisomal protrusions and tubules (Binns et al., 2006; Gao and Goodman, 2015; Hayashi et al., 2001; Schrader, 2001; Schuldiner and Bohnert, 2017). Peroxisomal extensions were also suggested to transfer the *A. thaliana* lipase SDP1 to lipid droplets during early seedling growth (Thazar-Poulot et al., 2015). In our hands, tubular peroxisomes were apparent during high overexpression of Far1 in COS-7 cells (not shown) but were only very rarely observed in U-2 OS cells.

Heterologous expression of human Far1 in the yeast *S. cerevisiae* indicated that the localization to LDs is not universally conserved. Far1 was localized to the ER in glucose medium, and to the vacuole when oleate was the main energy and carbon source (Fig. S1), but neither to peroxisomes nor to lipid droplets. Since sorting of *mammalian* Far1 to peroxisomes is pex19 dependent (Honsho et al., 2013), this suggests that the yeast pex19 homologue does not recognize the sorting motif contained within human Far1.

Targeting of proteins to lipid droplets may be broadly categorized by their respective original localizations, i.e. the ER or the cytosol (Kory et al., 2016; Ohsaki et al., 2014; Thiele and Spandl, 2008). ER derived membrane proteins appear to lack intralumenal domains so that they may be readily accommodated in the monolayer LD membrane; they also revert to the ER if lipid droplets are consumed during starvation (Zehmer et al., 2009). Far1 does not fit well into this class of LD proteins because it is neither an ER protein originally nor is it located at the ER under starvation. It also displays an intralumenal domain while on peroxisomes (Honsho et al., 2013) and is partially located at the ER (as evidenced by N-glycosylation; Fig. 2).

Integral transmembrane domains and the corresponding topology are usually well predicted by the TMHMM algorithm (Krogh et al., 2001). However, when it comes to lipid droplet associated proteins we found it rarely to be consistent with experimental data (not shown), presumably because of the obvious difficulty of squeezing a "standard" TMD into the phospholipid *monolayer*. Hairpin topologies and amphipathic helices, which are typical for LD proteins pose a formidable future challenge to bioinformatics.

### The targeting mechanism of Far1

The role of the ER for peroxisomal targeting and biogenesis is discussed controversially (Buentzel et al., 2015; Erdmann, 2016; Lodhi and Semenkovich, 2014; Mayerhofer, 2016; Smith and Aitchison, 2013). N-glycosylation of the Far1 reporter proteins (Fig. 2) indicated that about 50% of Far1 associated transiently with the ER in our model system, presumably *before* further transport towards the peroxisome was accomplished.

Pex19 is a cytosolic chaperone implicated in the targeting of peroxisomal membrane proteins (Jones et al., 2004), and it is also required for the sorting of Far1 to peroxisomes (Honsho et al., 2013). This traditional view has been extended by recent findings that pex19 is also involved in transporting the hairpin protein UBXD8 to the *endoplasmic reticulum*, from where it moves onwards to lipid droplets (Schrul and Kopito, 2016). However, while our pex19-KO cells replicated the expected phenotype (loss of peroxisomes), the localization of Far1 on lipid droplets or at the ER was not impaired (Fig. 5AB).

Far1 is also a tail-anchored membrane protein (Honsho et al., 2013), which is at least transiently exposed to the N-glycosylation machinery at the ER (Fig. 2). Several chaperone assisted targeting pathways have been described (Borgese and Fasana, 2011; Guna et al., 2018). The probably most prominent sorting mechanism involves TRC40/Get3p/ASNA1, which was also required for targeting peroxisomal proteins to the ER (van der Zand et al., 2010). However, depletion of TRC40 did not affect Far1 LD localization to any significant extent (Fig. 5CD). In hindsight, our most likely explanation would be that the targeting of tail anchored proteins involves a rather redundant machinery (Rabu et al., 2009). We also tried a double pex19/ASNA1-KO approach, but again Far1 was still found on LDs (Fig. 5E). Knocking out even more chaperones would be expected to show severe side effects (the translocon component sec61ß is also a tail anchored protein), and was not tried here. We conclude that Far1 targeting to the ER and lipid droplets is robustly supported by the cellular machinery of mammalian cells and is unlikely to be dependent on one single factor.

### Far1 is a novel dual topology membrane protein

We propose that Far1 may be integrated into the ER membrane in two different ways (Fig. 6). The first topology features an integral transmembrane domain (first hydrophobic domain; HD1) and an amphipathic helix-like second domain (HD2), which associates primarily with the lumenal leaflet of the ER membrane. This also appears to be the only form competent for export/integration to peroxisomes and satisfyingly explains our glycosylation and permeabilization data. The alternative second topology might be driven by a strong preference of the HD2 domain for the cytosolic leaflet of the ER, especially when triglycerides start to accumulate after fatty acid supplementation. Molecular dynamics simulation indicates that a 4 nm thick neutral lipid layer would already change the properties of the surrounding phospholipid monolayer significantly (Prevost et al., 2018), long before the budding of a LD would occur. If HD2 associates first with the membrane, the short loop of intermittent amino acids would no longer allow an integral transmembrane orientation for HD1. This topology has the short C-terminal region exposed to the cytosol and is therefore in line with our permeabilization data for LD localized Far1. HD2 appeared as efficient as the native ACSL3 hydrophobic domain in terms of LD targeting (Fig. 4D). ACSL3 is considered a bona fide lipid droplet protein (Fujimoto et al., 2004; Poppelreuther et al., 2012; Walther et al., 2017) and is already present on emerging LDs (Kassan et al., 2013).

**Figure 6:**
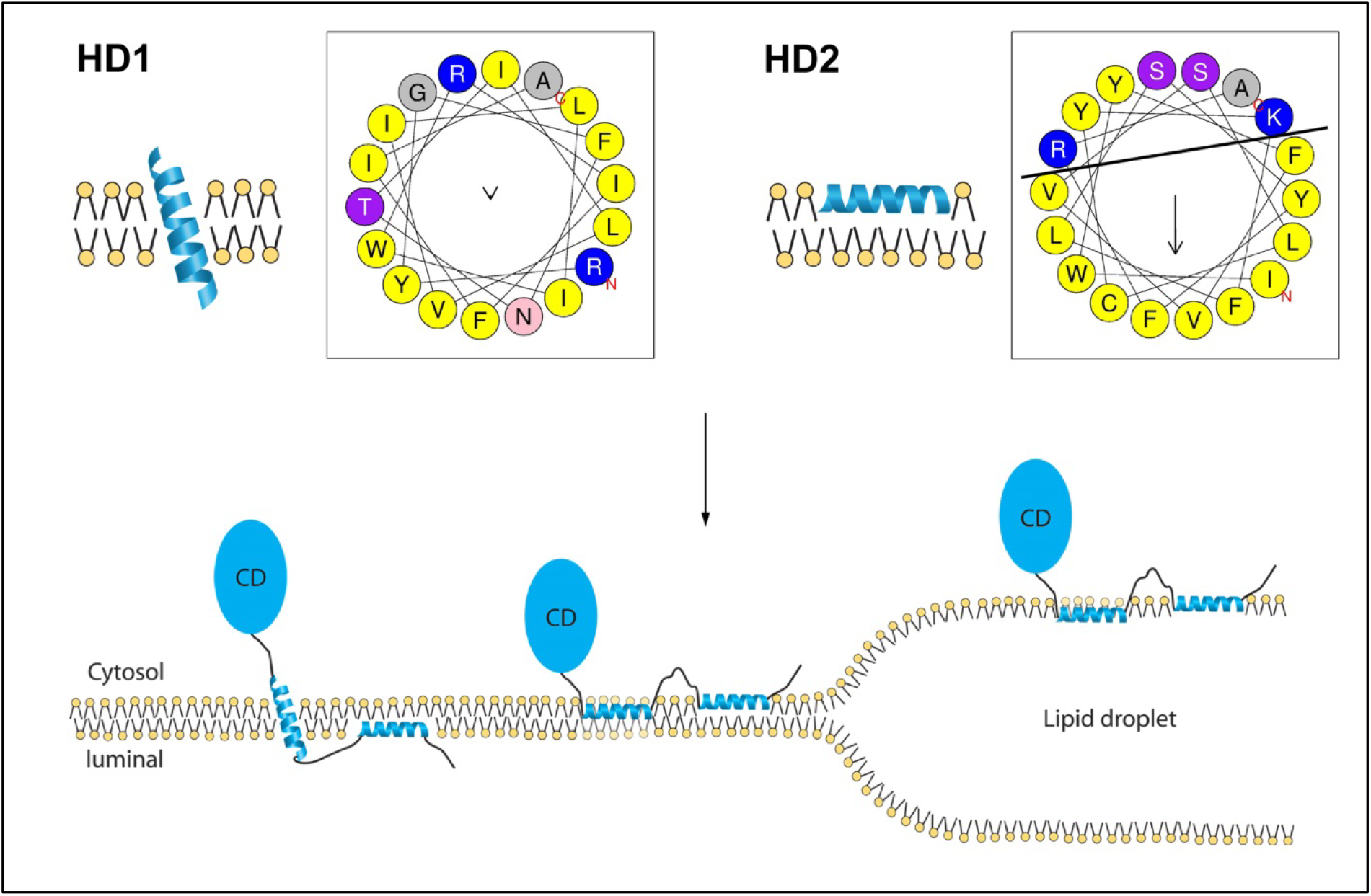
Topologies of the two hydrophobic domains and Far1. The first hydrophobic domain (HD1) most likely assumes an integral TMD configuration (Figs. 3D, 4A), whereas the second HD2 forms either an amphipathic helix as depicted here (Figs. 3C, 4C), or a hairpin with the turn embedded in the membrane bilayer. Helical wheel projections of the hydrophobic domains HD1 (I466-A484) and HD2 (I492-A509) show that the second hydrophobic domain displays a remarkable sidedness (black line), with charged and polar amino acids clustered to one side. Two hypothetical alternative topologies of Far1 are shown below. On peroxisomes, Far1 very likely assumes the leftmost configuration, with the C-terminus oriented towards the lumen. On lipid droplets, the neutral lipid core expels all charged amino acids towards the surrounding phospholipid monolayer. At the ER, Far1 presumably exists in two alternative conformations, with the leftmost topology consistent with the observed glycosylation tagging (Fig. 2B) and the ER being a transient trafficking itinerary towards peroxisomes. The alternative topology depicted in the middle would allow smooth transfer to lipid droplets after their induction by oleate supplementation. HD: hydrophobic domain; CD: catalytic domain.

Alternative scenarios may involve direct trafficking from peroxisomes to LDs as exemplified by the *A. thaliana* lipase SDP1 (Thazar-Poulot et al., 2015) or a retrograde transport from peroxisomes to the ER (Passreiter, 1998; Titorenko and Rachubinski, 1998). However this would involve moving the lumenal short hydrophilic segments of the C-terminus across the hydrophobic phospholipid layer which is considered energetically highly unfavorable.

Dual topology is a rare phenomenon, and most cases described so far involve multipass transmembrane domain proteins (>3 TMDs; Lambert and Prange, 2001; Rapp et al., 2006; Stockbridge et al., 2014; Wurie et al., 2011). Generally, the orientation of membrane proteins is considered to be determined during their cotranslational insertion into the membrane, but post-assembly topology switching does occur (Bogdanov et al., 2014). We applied the term "dual topology" to mean that one amino acid sequence occurs in two different topological orientations with respect to a phospholipid membrane (Granseth, 2010). However, dual topology has also been more narrowly defined as proteins which "insert into the membrane in two opposite orientations with an approximate 1:1 stoichiometry" (von Heijne, 2006). This would not encompass Far1, because the alternative topology of Far1 is restricted to the C-terminus and does not change the cytosolic orientation of the enzyme domain. Moreover, the stoichiometry is not fixed but dependent on the extent of triglyceride synthesis.

The topologies of Far1 appear to be strongly correlated to the subcellular localization. Far1 converts fatty acyl-CoA to fatty alcohols; on peroxisomes these are further metabolized by lumenal enzymes and are finally incorporated into ether lipids. Far1 on lipid droplets may therefore reduce the production of ether lipids from exogenously supplied fatty acids. The metabolic fate of LD synthesized fatty alcohols is currently unclear and beyond the scope of this study. They could enter a fatty alcohol cycle, which would be involved in cellular lipid homeostasis (Rizzo, 2014), balancing ether and non-ether lipid levels. In line with this, the vast majority of externally added hexadecanol was converted to palmitate rather than to ether lipids in cultured fibroblasts (Rizzo et al., 1987). Other metabolic routes are also conceivable, as many enzymes of lipid metabolism are found on LDs (Goodman, 2009).

Finally, preliminary but tantalizing recent data suggest that lipid droplet and peroxisome biogenesis may occur at the same ER subdomains marked by mammalian MCTP2/yeast Pex30 and seipin (Joshi et al., 2017). This spatial proximity may well be the mechanistic basis for the alternative sorting pathways of Far1, and may also be instrumental in allowing dual topologies upon insertion into the membrane.

## Acknowledgements

We thank the Electron Microscopy Core Facility (EMCF) at Heidelberg University, especially Uta Haselmann and Simone Hoppe for their technical assistance with the correlative-light-electron-microscopy. Furthermore, we would like to thank Dr. Vibor Laketa and the Infectious Diseases Imaging Platform (DIP; Department for Infectious Diseases) for microscopy support. Juan Liao, Simone Sander and Simon Gairing from our group contributed plasmids for this study. T.E. was supported by the MD/PhD program of the Medical Faculty of the University of Heidelberg.

Data were discussed with Blanche Schwappach (University of Göttingen), and J. R.-M. was funded by SFB 1002 (TP A07 to B.S.). Funding by the German research foundation DFG (FU 340/7-1 to J.F.), and the Stiftung Nephrologie Heidelberg (to M.P. and J.F.) is gratefully acknowledged. R.B. was supported by the Deutsche Forschungsgemeinschaft (TRR83, TP13).

## Supplement

**Suppl. Figure S1.**
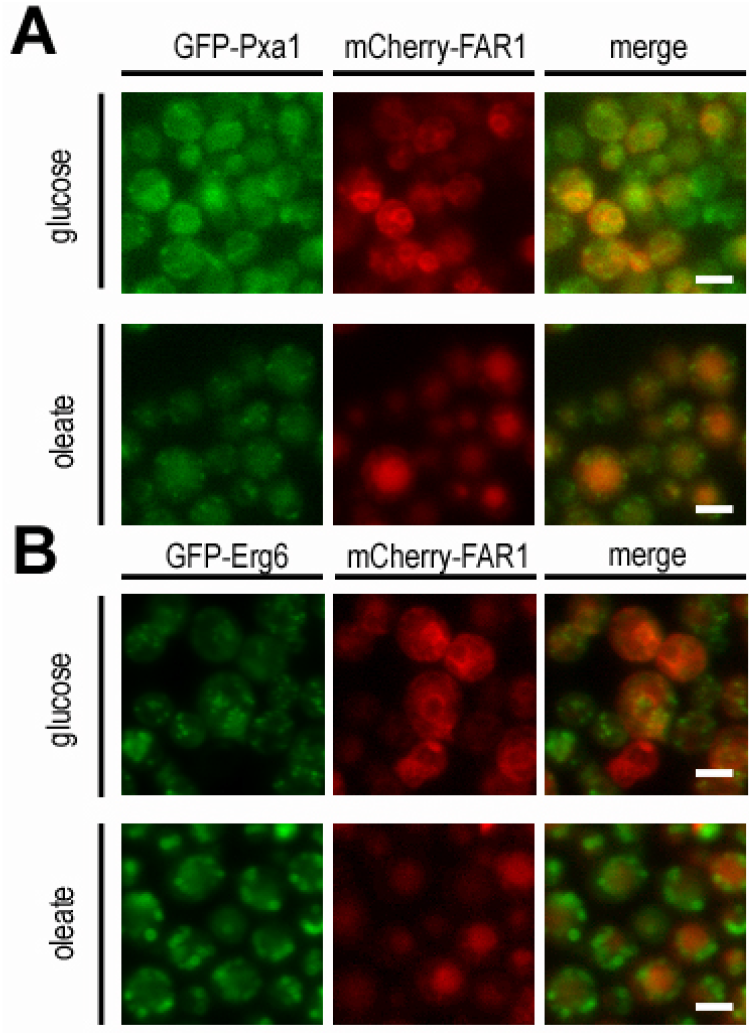
Human Far1 does not localize to peroxisomes and lipid droplets in yeast. (A) Far1 does not overlap with the peroxisomal marker GFP-Pxa1 in yeast. Yeast cells were transformed with the cDNA coding for the human Far1 (red). No overlap can be seen with endogenous GFP-Pxa1 under glucose or oleate conditions. Instead, Far1 is localized to the ER (glucose) or in vacuoles (oleate). (B) Far1 does not overlap with the lipid droplet marker Erg6 in yeast. Yeast cells were transformed with mCherry-Far1. No overlap can be seen with lipid droplets. Instead, Far1 is localized to the ER (glucose) or in vacuoles (oleate).

**Suppl. Figure S2.**
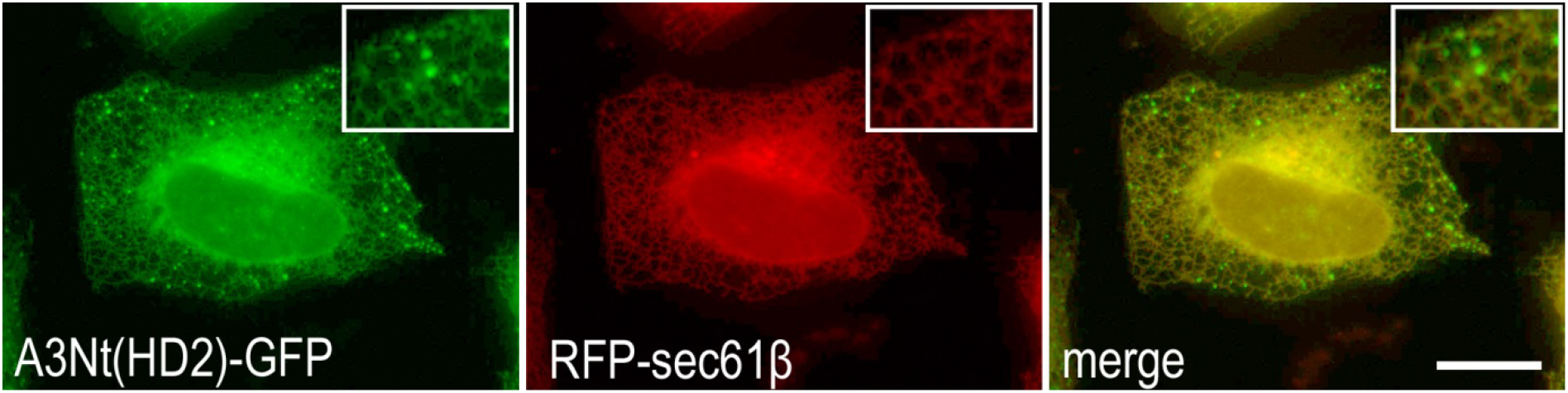
The second hydrophobic domain can substitute for the amphipathic helix of ACSL3. The amphipathic helix of ACSL3-N-terminus was exchanged for the second hydrophobic domain of Far1 and expressed in U-2 OS cells. Under steady state conditions, the signal derived from GFP overlaps convincingly with the ER marker sec61ß. Additional small green puncta (inset) correspond to small lipid droplets that are only stained by A3Nt(HD2)-GFP. Standard growth medium (-OA). Scale bar 10μm

**Suppl. Figure S3.**
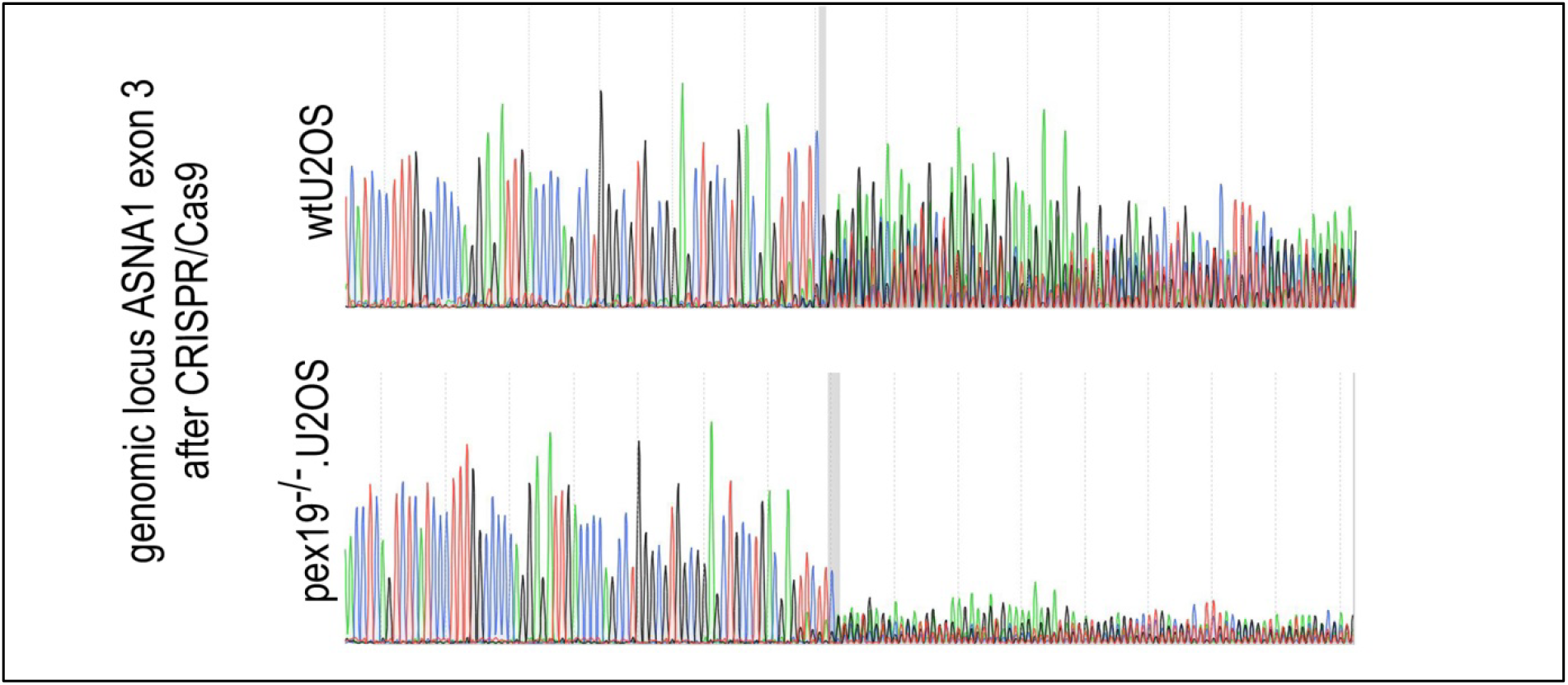
Sequencing of the genomic locus of ASNA1 exon 3 suggests highly decomposed sequence. Knockout pool cells were generated as described in the materials and methods part and harvested. The genomic locus of the expected Cas9 cutting site was amplified and sequenced. The grey bar indicates the expected cutting site of Cas9. Decomposed sequence is visible behind the predicted cutting site.

## Abbreviations

Far1: fatty acyl CoA reductase 1
Far1Ct: Far1 C-terminus
LD: lipid droplet
ER: endoplasmic reticulum
HA: hemagglutinin epitope
A3Nt: ACSL3 N-terminus
GFP: green fluorescent protein
FATP4: fatty acid transport protein 4
OA: oleic acid
HD: hydrophobic domain

